# An infectious Rous Sarcoma Virus Gag mutant that is defective in nuclear cycling

**DOI:** 10.1101/2021.04.16.440250

**Authors:** Clifton L Ricaña, Marc C Johnson

## Abstract

During retroviral replication, unspliced viral genomic RNA (gRNA) must escape the nucleus for translation into viral proteins and packaging into virions. “Complex” retroviruses such as Human Immunodeficiency Virus (HIV) use cis-acting elements on the unspliced gRNA in conjunction with trans-acting viral proteins to facilitate this escape. “Simple” retroviruses such as Mason-Pfizer Monkey Virus (MPMV) and Murine Leukemia Virus (MLV) exclusively use cis-acting elements on the gRNA in conjunction with host nuclear export proteins for nuclear escape. Uniquely, the simple retrovirus Rous Sarcoma Virus (RSV) has a Gag structural protein that cycles through the nucleus prior to plasma membrane binding. This trafficking has been implicated in facilitating gRNA nuclear export and is thought to be a required mechanism. Previously described mutants that abolish nuclear cycling displayed enhanced plasma membrane binding, enhanced virion release, and a significant loss in genome incorporation resulting in loss of infectivity. Here, we describe a nuclear cycling deficient RSV Gag mutant that has similar plasma membrane binding and genome incorporation to WT virus and surprisingly, is replication competent albeit with a slower rate of spread compared to WT. This mutant suggests that RSV Gag nuclear cycling is not strictly required for RSV replication.

**Importance:** While mechanisms for retroviral Gag assembly at the plasma membrane are beginning to be characterized, characterization of intermediate trafficking locales remain elusive. This is in part due to the difficulty of tracking individual proteins from translation to plasma membrane binding. RSV Gag nuclear cycling is a unique phenotype that may provide comparative insight to viral trafficking evolution and may present a model intermediate to cis- and trans-acting mechanisms for gRNA export.

## Introduction

Retroviruses hijack a multitude of host processes to overcome barriers throughout the viral lifecycle. One such barrier is the nuclear membrane, which protects host genetic data and allows for regulation of genes by keeping unspliced RNA transcripts from exiting the nucleus. Retroviral genomic RNA (gRNA) consists of a long unspliced transcript that must escape the nucleus and traffic to the plasma membrane for virion packaging. Retroviruses, such as HIV, have evolved trans-acting viral proteins to facilitate the active transport of unspliced gRNA out of the nucleus via interaction with cis-acting elements on the gRNA. An Arg-rich nuclear localization signal (NLS) on the Rev protein of HIV allows nuclear entry of non-gRNA-bound Rev via importin-β (1, 2). A nuclear export signal (NES) on Rev allows nuclear export of gRNA-bound Rev via the exportin Chromosomal Maintenance 1 (CRM1) (1, 3, 4). “Simpler” retroviruses such as Mason-Pfizer Monkey Virus (MPMV) and Murine Leukemia Virus (MLV) exclusively use gRNA cis-acting elements in conjunction with host cell export factors (5–8). In the case of the alpharetrovirus, Rous Sarcoma Virus (RSV), evidence has implicated the Gag structural protein nuclear cycling as a trans-acting mechanism for exporting gRNA (reviewed in (9–11)).

Initial gross truncation of RSV to study plasma membrane binding unexpectedly found that a Matrix (MA)-GFP fusion protein was enriched in the nucleus (12). Since RSV-Gag-GFP with Protease (PR) deleted (hereafter referred to as RSV-Gag^WT^) expresses in the cytoplasm/plasma membrane at steady state, this pointed toward the full Gag protein potentially trafficking to intermediate subcellular locales (12). To determine whether the non-nuclear phenotype of Gag was due to size exclusion or nuclear export, Leptomycin B (LMB) was used to block the CRM1 export pathway of cells transfected with RSV-Gag^WT^ (12). With LMB treatment, RSV-Gag^WT^ was shown to rapidly shift to an almost exclusive nuclear expression (12). Truncation of RSV-Gag^WT^ and amino acid manipulation demonstrated that the NES was located in the p10 domain and that a single (L219A) point mutation blocked nuclear export resulting in only nuclear expression (12, 13). Further characterization verified NLSs consisting of a non-canonical importin-11 and transportin-3 (TNPO3) dependent NLS in the tertiary structure of MA and a canonical four basic amino acid importin-α/β dependent NLS motif in nucleocapsid (NC, K_36_KRK_39_) (12, 14–17).

Interestingly, a non-infectious mutant Myr1E that did not accumulate in the nucleus with LMB treatment exhibited strong plasma membrane binding, increased virion release (1.4xWild Type, WT), and had a defect in genome packaging (0.4xWT) (12). This suggested a Gag nuclear localization requirement for genome packaging (12, 18). Myr1E consists of the myristoylated 10-amino acid Src plasma membrane binding domain added to the N-terminal end of RSV-Gag (12, 19). Another non-infectious mutant, SuperM, consists of RSV-Gag with two Glu swapped with Lys (E25K and E70K) (20). SuperM also displayed strong plasma membrane binding that did not accumulate in the nucleus with LMB treatment, increased viral release (3xWT), and had a severe defect in genome packaging (0.1xWT) (20). To characterize the role of Gag nuclear localization in genome packaging, a canonical NLS was engineered into Myr1E.NLS (17, 18). Though both viruses were not infectious; nuclear cycling was enhanced with Myr1E.NLS as compared to Myr1E, viral release remained the same (1.4xWT), and corresponded to recovered genome packaging at nearly WT levels (18). To complement this finding, a different study showed that Gag exhibited reduced binding to nuclear import factors importin-α and -11 when bound to viral RNA containing the Ψ packaging signal (21). In conjunction, Gag bound to ΨRNA promoted binding to CRM1 (21). Together, these data suggest that RSV requires Gag nuclear cycling to export unspliced gRNA for subsequent packaging and infectivity.

While the previously summarized data suggests Gag involvement with gRNA nuclear export and packaging, there is also evidence against this argument. There is a gRNA cis-acting element found in the direct repeats (DR) flanking the *Src* gene of RSV (22, 23). DR deletion and various DR mutants severely decreased the amount of unspliced RNA transcripts and Gag protein levels in the cytoplasm of chicken embryo fibroblasts (22, 23). In a gain of function assay, expression of HIV-1 Gag protein, a proxy for unspliced HIV-1 gRNA translocation into the cytoplasm, was recovered when RSV DRs were engineered into HIV-1 constructs devoid of functional Rev protein and Rev response element on the gRNA (22). To parse the mechanism of nuclear export, the investigators used a three-part assay consisting of RSV constructs encoding chloramphenicol acetyl transferase (CAT) to report unspliced RNA in the cytoplasm of transfected cells, fluorescent in situ hybridization (FISH) of reporter viral RNA (vRNA) containing DR, and dominant-negative mutants of host cellular nuclear export factors Tap and Dbp5. With this assay, the DR were shown to use and require Tap and Dbp5 to translocate the viral RNA to the cytoplasm (24). Furthermore, addition of increasing amounts of transfected RSV Gag did not increase cytoplasmic levels of DR containing CAT reporter vRNA nor did it increase cytoplasmic levels of Ψ containing CAT reporter vRNA (24). This line of evidence is corroborated by other studies and mutational analyses which point toward both DR playing a role in Gag assembly and DR2 as the major cis-acting element for unspliced RNA transport (25–27). These pieces of evidence suggest that RSV Gag nuclear cycling is not involved with or plays a minor role in gRNA translocation to the cytoplasm.

In a survey of various retroviral Gag proteins, our laboratory described a non-nuclear cycling HIV-MA/RSV-Gag-GFP^ΔPR^ (H/RSV-Gag) chimera. Cloning this chimera into a single-cycle H/RSV^ΔEnv^ provirus demonstrated that it produced about half as many infectious particles as WT virus (28). HIV-MA is myristoylated like the non-infectious Myr1E RSV mutant and presumably would cause similar genome packaging defects that result in severe infectivity defects, but this is not the case and suggests additional mechanisms are involved. Alteration of plasma membrane trafficking and membrane binding has consequences not limited to genome packaging (29–32). Briefly, non-myristoylated Gag-membrane interactions are mainly due to electrostatic interactions and altered hydrophobic interactions with acyl chains such as phosphatidylinositol-4,5-bisphosphate (PI(4,5)P_2_) or cholesterol depletion can reduce plasma membrane binding (29–32). Separating the effects of plasma membrane binding and/or trafficking alterations from nuclear cycling would provide fewer confounding interpretations of nuclear cycling effects on viral assembly. In addition, the separation of the functions in a replication competent context would provide insight into overall effects of nuclear cycling during the viral lifecycle. Replication competency of our infectious, non-nuclear cycling H/RSV chimera, however, could not be tested due to the major splice donor for Env located in the N-terminus of MA (28, 33). Here, we characterize a replication competent mutant that contains a minimal amino acid substitution from a polymorphism found in the NC-NLS of RSV strain JS11C1, is defective in nuclear cycling, and incorporates gRNA at similar levels to WT.

## Results

### Validation of NLS’s and NES in RSV-Gag and determination of the H/RSV-Gag^WT^ nuclear cycling block

We demonstrated previously that replacement of the RSV MA domain with HIV-1 MA prevents nuclear cycling of RSV Gag. To probe how HIV-MA blocks nuclear cycling in the H/RSV-Gag^WT^ construct, we tested various RSV deletion and HIV-MA mutation/truncation constructs in the RSV-Gag^WT^ and H/RSV-Gag^WT^ backbone respectively (Fig 1A). Plasmid constructs were transfected in parallel into DF1 cells (chicken embryonic fibroblast) and imaged 16-18 hrs post transfection followed by LMB (10 ng/mL) treatment for 1 hr and further imaging. At steady state, RSV-Gag^WT^ and H/RSV-Gag^WT^ displayed diffuse and in many instances both diffuse and punctate cytoplasmic expression of GFP in contrast to the nearly exclusive nuclear expression of the previously described RSV-Gag^L219A^ mutant (Fig 1B) (13, 28). Recapitulating previous findings, CRM1 inhibition with LMB resulted in accumulation of RSV-Gag^WT^ to the nucleus but not H/RSV-Gag^WT^ (Fig 1C) (13, 28). To further validate our system, previously described mutants, RSV-Gag^ΔMA^ and RSV-Gag^ΔNC^ (14), in addition to RSV-CANC were tested for nuclear accumulation (Fig 1B and C). As expected, RSV-Gag^ΔMA^ displayed nuclear accumulation with LMB treatment indicating strong nuclear import via the NC-NLS, followed by strong export of the Gag protein via the p10-NES. Recapitulating previous findings, RSV-Gag^ΔNC^ expressed in relatively equal amounts in the cytoplasm and nucleus with LMB treatment (14). This suggests that the NC-NLS is the dominant NLS in RSV Gag. RSV-CANC, lacking the NES in p10, displayed nearly exclusive nuclear expression with and without LMB (Fig 1B and C). This indicates the NC-NLS is sufficient to drive nuclear import and that HIV-MA is somehow able to inhibit nuclear import in H/RSV-Gag^WT^.

**Figure 1.**
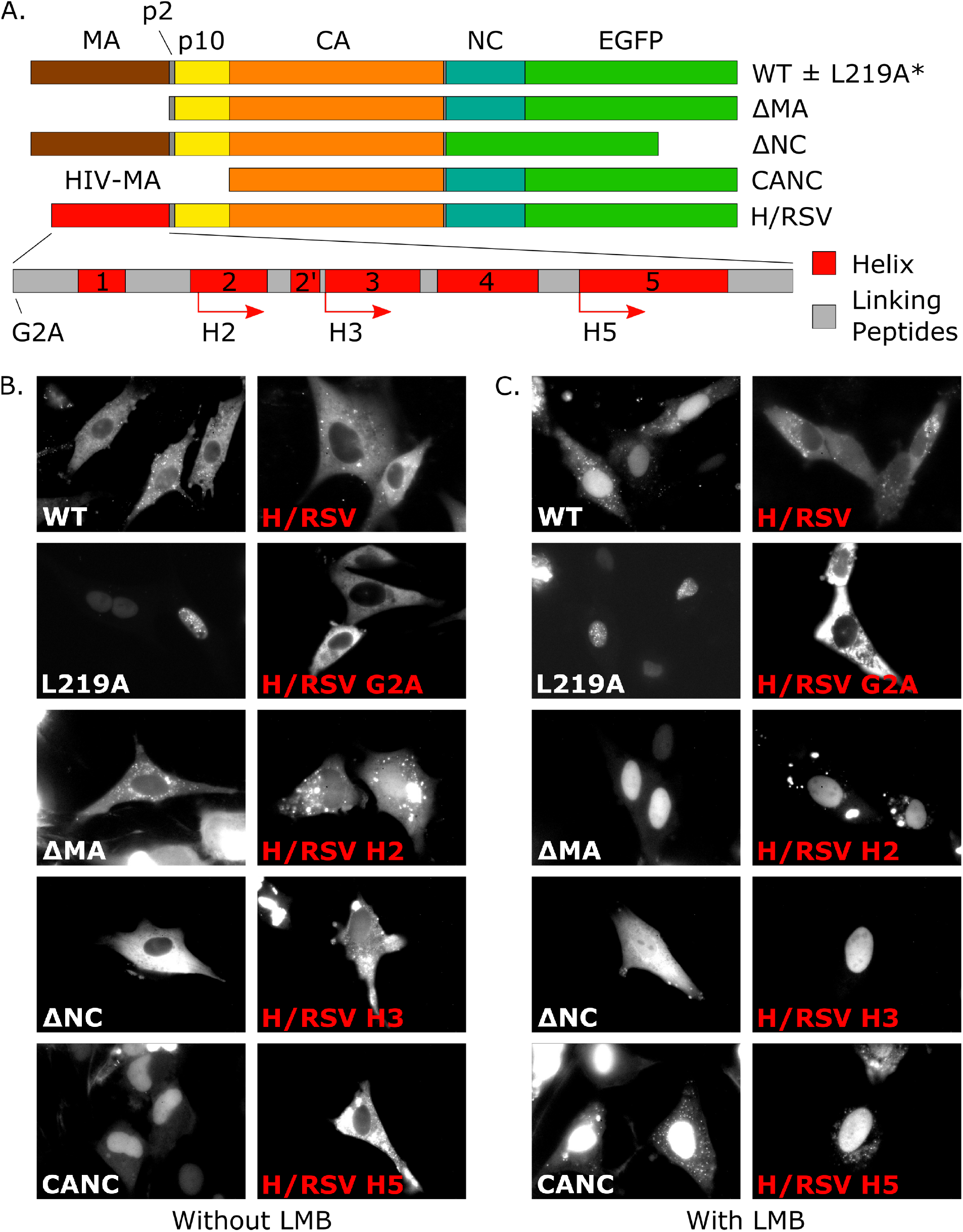
Subcellular expression of various Gag-GFP fusion proteins. (A) Schematic of Gag-GFP fusion proteins tested. RSV^WT^, RSV^L219A^ (lacks a functional NES), RSV^ΔMA^ (removes one NLS), RSV^ΔNC^ (removes another NLS), and RSV-CANC were tested in parallel. H/RSV-Gag^WT^ was further probed with the G2A mutation (blocks myristoylation thus plasma membrane targeting) and α-helix truncations from HIV-MA. α-Helices are labeled in red and linking peptides, consisting of strands and loops, are in grey. For α-helix truncations, the starting Met was kept followed by the α-helix of interest. H/RSV-Gag^H2^ truncates helix-1 through the first four amino acids of helix-2 (K30-H33) and contains the remainder of helix-2 through the end of HIV-MA (I34-Y132). H/RSV^H3^ truncates helix-1, helix-2, helix-2’, and begins at the N-terminus of helix-3 (T55-Y132). H/RSV-Gag^H5^ truncates helix-1 through -4 and begins at the N-terminus of helix-5 (D96-Y132). (B-C) Representative images of steady state expression of Gag-GFP fusion proteins in DF1 cells. (B) Cells were imaged starting 16-18 hrs post transfection and (C) imaged after LMB (10 ng/mL) treatment for 1 hr.

HIV Gag, unlike RSV, is myristoylated and this modification is required for proper HIV Gag trafficking and binding to the plasma membrane (30, 34). To probe whether myristoylation and plasma membrane trafficking potentially counteract the NLS signal, a known myristoylation defective G2A mutation was tested in the H/RSV-Gag^WT^ construct (35). H/RSV-Gag^G2A^ expressed primarily in the cytoplasm with little to no plasma membrane accumulation and no residual puncta on the cell culture dish that would presumably be budded H/RSV-Gag^G2A^ virus like particles (VLPs) (Fig 1B). Additionally, no change in phenotype was seen with LMB treatment (Fig 1C). This suggests that membrane binding via PI(4,5)P_2_ interaction with myristoylated Gag is not the determining factor for the blocked nuclear import of H/RSV-Gag^WT^. A possible explanation is that HIV-MA sterically hinders the NC-NLS, or alternatively, HIV-MA counteracts the NC-NLS by anchoring the polyprotein to some feature in the cytoplasm (30).

To locate HIV-MA residues important for the cytoplasmic retention of H/RSV-Gag^WT^, we used previously characterized secondary structural domains to sequentially truncate α-helices from the N-terminus of the H/RSV-Gag^WT^ backbone, leaving the starting Met followed by the α-helix of interest (36). H/RSV-Gag^H2^ truncates helix-1 through the first four amino acids of helix-2 (K30-H33) and contains the remainder of helix-2 through the end of HIV-MA (I34-Y132) (Fig 1A). H/RSV-Gag^H3^ truncates helix-1, helix-2, helix-2’, and begins at the N-terminus of helix-3 (T55-Y132) (Fig 1A). H/RSV-Gag^H5^ truncates helix-1 through -4 and begins at the N-terminus of helix-5 (D96-Y132) (Fig 1A). All three of the truncations of HIV-1 MA restored nuclear cycling of H/RSV-Gag^WT^ (Fig 1B-C). Interestingly, H/RSV-Gag^H2^ and H/RSV-Gag^H3^ displayed varying degrees of nuclear expression even in the absence of LMB (Fig 1B). These data demonstrate that the nuclear retention H/RSV-Gag^WT^ requires the presence of helix-1 of HIV-1 MA.

### An alternative Gag mutant separating plasma membrane trafficking and nuclear cycling

Non-nuclear cycling H/RSV^ΔEnv^ pseudotypes were infectious, causing us to question the importance and requirement of nuclear cycling during the RSV lifecycle. Previously described mutants Myr1E and SuperM were presumed to overcome nuclear trafficking by enhancing targeting to the plasma membrane resulting in the loss of genome incorporation (12, 18, 20). However, the loss may also result from too rapid of virion escape excluding genome incorporation. Parsing the mechanism requires separation of plasma membrane trafficking from nuclear localization. To further probe the dominant NC-NLS, we performed a BLAST database search for known polymorphisms at the characterized NC-NLS (15). We found one polymorphism, NC-K36E, in the infectious clone (JS11C1) that was constructed from the genomic viral sequence isolated from a chicken breed in a study by Cui and colleagues (37).

In the previous study by Lochmann and colleagues, the NC-NLS was removed by changing all four positive charges to Ala (15). This Ala mutant represents a fairly significant change in peptide composition and may have had unintended phenotypic consequences. Since the study that identified JS11C1 was focused on general characterization and comparison of avian leukosis viruses prevalent in indigenous agricultural chickens, specific amino acid polymorphisms were probably not as carefully scrutinized and single, isolated divergent amino acids may have arisen from sequencing artifacts. To test the phenotype of the polymorphism in JS11C1, the NC-K36E mutation was engineered into RSV-Gag^K36E^ (Fig 2A). Because NC-K36E is an opposite charge change, we also engineered the more neutral and more structurally similar NC-K36M into RSV-Gag^K36M^. At steady state, RSV-Gag^WT^, RSV-Gag^K36E^, and RSV-Gag^K36M^ displayed diffuse and in many instances both diffuse and punctate cytoplasmic expression of GFP (Fig 2B). LMB treatment resulted in nuclear accumulation of RSV-Gag^WT^, but both RSV-Gag^K36E^ and RSV-Gag^K36M^ remained cytoplasmic even after one hour of treatment (Fig 2C). These data demonstrated that the single point mutation found in JS11C1 NC was sufficient to abolish the nuclear cycling of RSV Gag.

**Figure 2.**
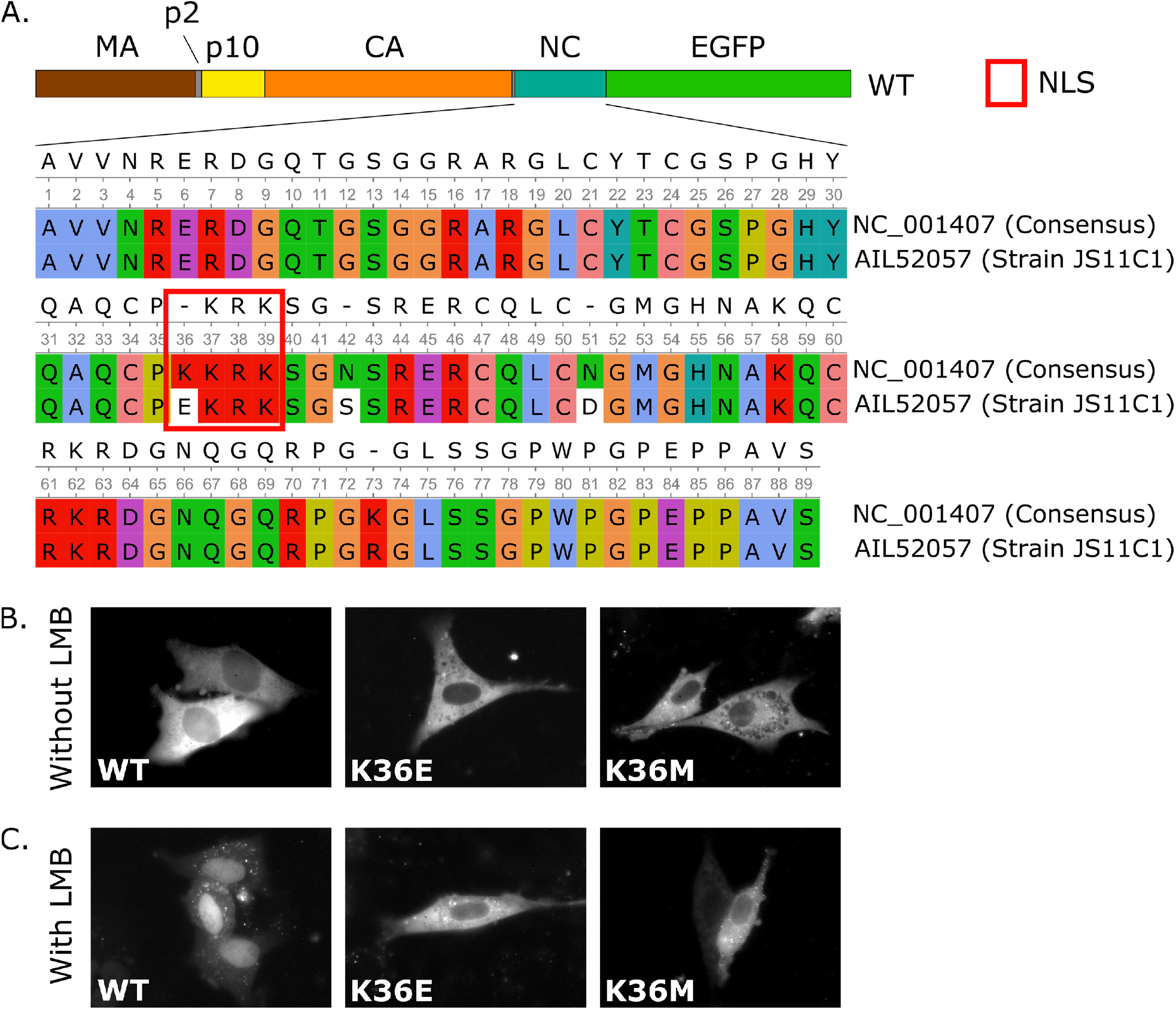
RSV-Gag^K36E/M^ mutants are defective in nuclear cycling. (A) Multiple sequence alignment of RSV-NC between the WT consensus and strain JS11C1. There is a K36E polymorphism in the NC-NLS. Since Glu is the opposite charge from Lys, both Glu and Met mutations were tested for nuclear import activity. Mutations were made in the WT backbone (panel A). (B) Representative images of steady state expression of RSV-Gag^WT^ and RSV-Gag^K36E/M^ EGFP fusion proteins in DF1 cells. Cells were imaged starting 16-18 hrs post transfection. (C) Representative images of expression of fusion proteins in cells from the previous panel after LMB (10 ng/mL) treatment for 1 hr.

### RSV^K36M^ is replication competent but has a reduced rate of spread compared to RSV^WT^

Compared to the RSV-Gag^WT^, NC-K36E/M mutations were deficient in nuclear cycling. To test whether these mutations were tolerated in infectious virus, we engineered NC-K36E/M into an RSV^ΔEnv^ single-cycle construct with two fluorescence reporters (Fig 3A). Briefly, crimson fluorescence protein is encoded outside of the viral genome, while EGFP is incorporated into the viral genome. This construct is psuedotyped with Vesicular Stomatitis Virus glycoprotein (VSV-g) upon transfection. Transfected cells fluoresce both crimson and green; however, subsequently infected, un-transfected target cells fluoresce only green since virions from the producer cell only encode for EGFP (Fig 3B). Distribution of fluorescent DF1 cells, as measured by the ratio of infected to transfected cells, was then determined via flow cytometry five days post transfection. RSV^ΔEnv,K36M^ and RSV^ΔEnv,K36M^ were able to infect target cells at about 50% and 10% the rate of RSV^ΔEnv^, respectively (N=6, p<0.01, pairwise Wilcoxson rank sum test, Fig 3C).

**Figure 3.**
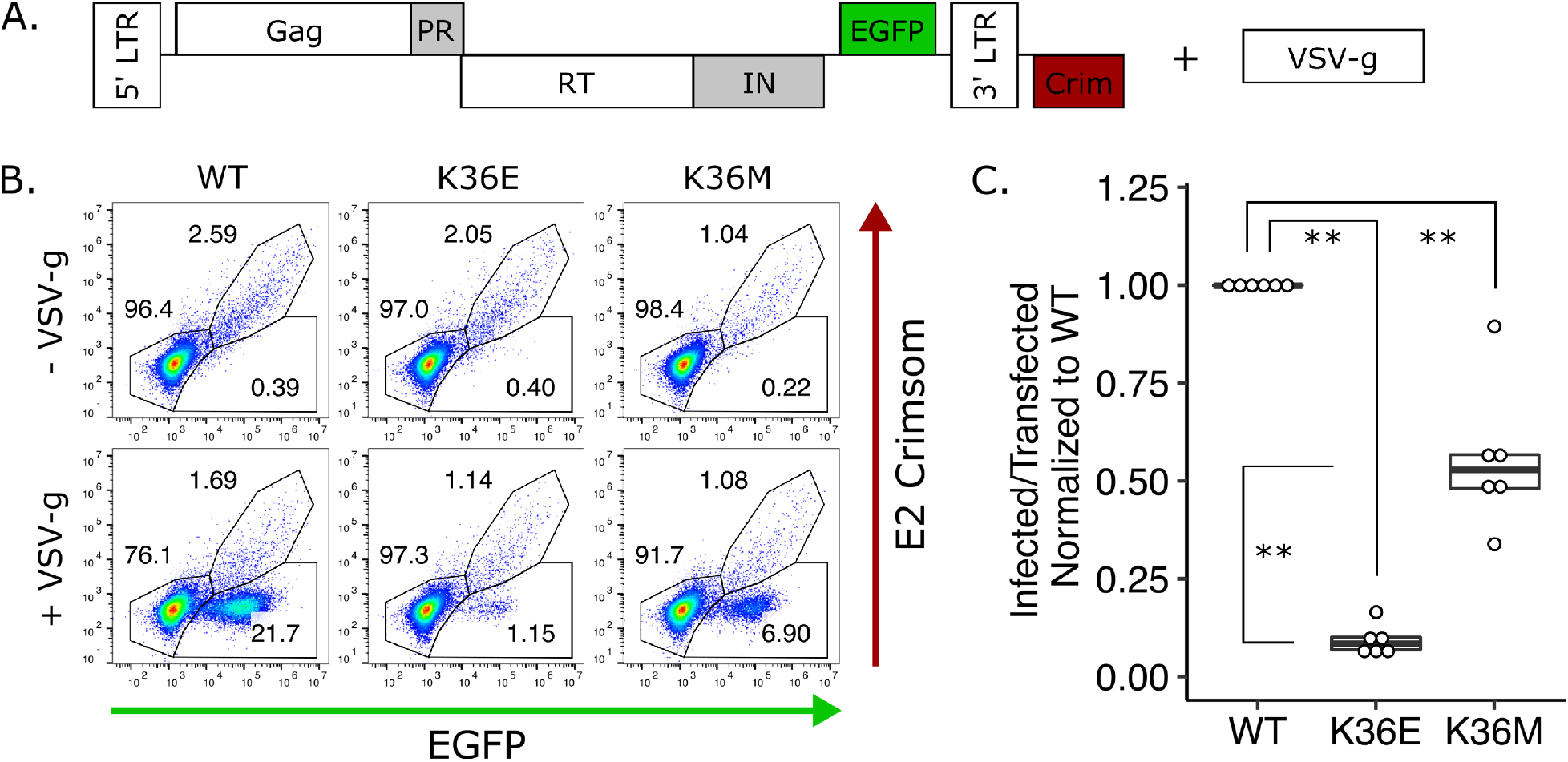
Gag NLS mutants are infectious in single-cycle virus. (A) Schematic of the single-cycle provirus tested (NC-K36E/M mutations not shown). Since the Crimson fluorescence gene is located outside of the viral genome (indicated by the LTRs), transfected cells are indicated by both GFP (+) and Crimson (+) cells, while subsequent infections are indicated by only GFP (+) cells. (B) Representative flow plots for single-cycle infection assays using a 2-color reporter system with and without VSV-g. Samples were collected five days post transfection for flow cytometry. (C) Ratio of infected to transfected cells normalized to WT. Statistics: N=6, ** p<0.01, pairwise Wilcoxson rank sum test.

To address the possibility of transfection or pseudotyping artifacts contributing to the apparent infectiousness of the mutants, NC-K36E/M were engineered into replication competent RCAS-GFP (RSV^WT^, Fig 4A). RSV^WT^, RSV^K36E^, and RSV^K36M^ were then transfected into separate dishes of DF1s. Both RSV^K36E^ and RSV^K36M^ were able to infect 75% or more cells within two weeks. To compare kinetics, the virus producing cell-line was co-cultured with uninfected target cells, starting at 1% of the total cell population, and a portion collected every other day for flow cytometry. Since RSV^ΔEnv,K36E^ displayed more severe defects in the single-cycle assay and RSV^K36E^ spread through tissue culture at a markedly slower rate than RSV^K36M^ in preliminary testing (data not shown), RSV^K36E^ was not further tested. RSV^K36M^ spread through tissue culture at a reduced rate compared to RSV^WT^. RSV^WT^ spread to 93% of the culture on day six, while RSV^K36M^ only spread to 72% on day six (N=5, p<0.01, pairwise Wilcoxson rank sum test, Fig 4B). Together, the slowed kinetics of replication competent RSV^K36M^ recapitulates the dysfunction seen in RSV^ΔEnv,K36M^.

**Figure 4.**
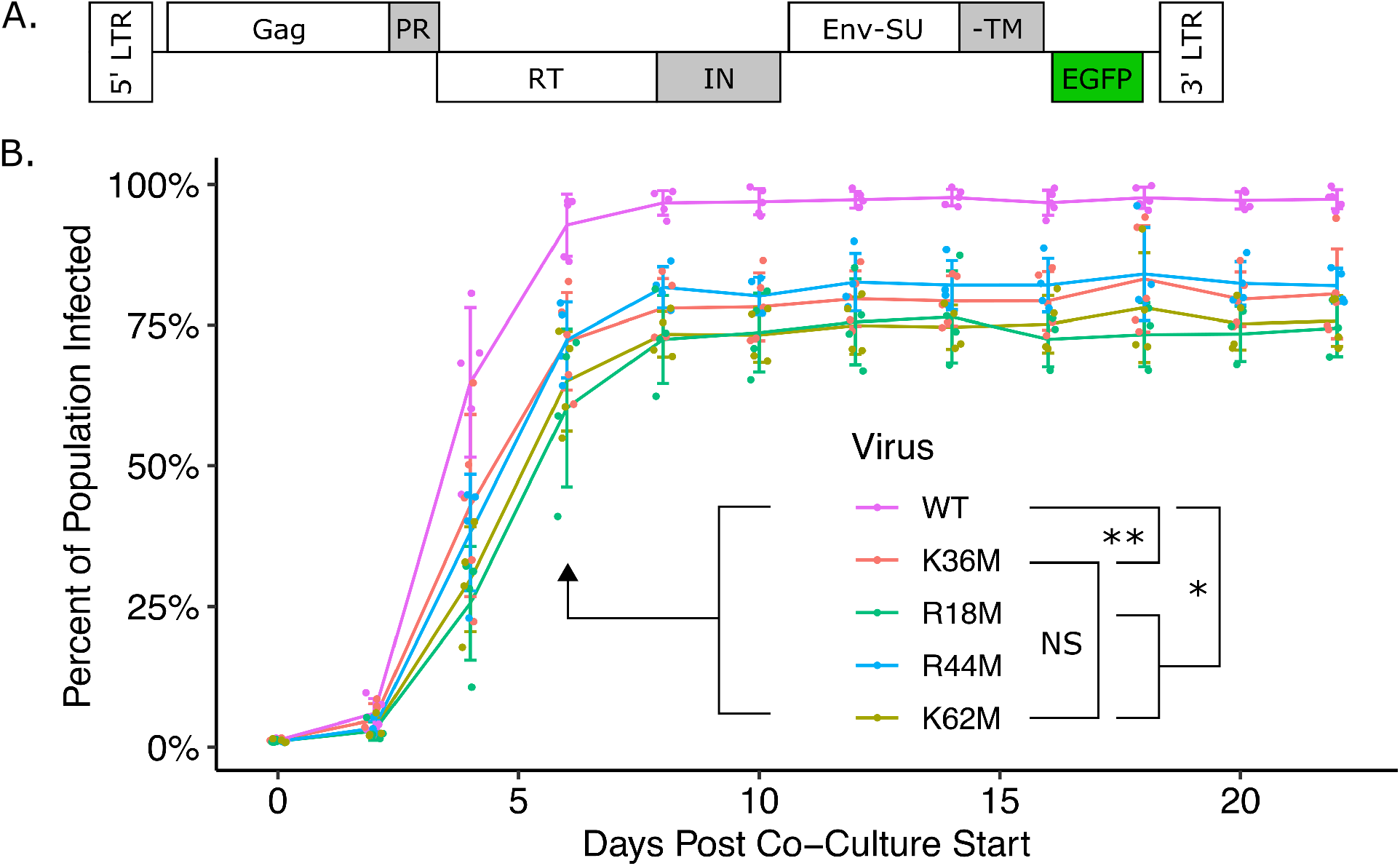
Replication competent RSV^K36M^ virus spreads albeit at a slower rate than WT. Since RSV^ΔEnv/K36M^ displayed near RSV^ΔEnv^ levels of spread compared to RSV^ΔEnv/K36E^ in Fig 3, experiments carried forward used RSV^K36M^ to reduce assay times. (A) Schematic of replication competent virus with an EGFP reporter in place of Src (K36M mutation not shown). (B) Replication competent virus spread through cell culture as measured by percent GFP (+) cells. Briefly, uninfected target DF1 cells were co-cultured with 1% fully infected cells. Samples were collected every other day for EGFP quantification via flow cytometry. Since positive amino acids in NC are important for nucleic acid binding, three other positive residues in NC were separately mutated to Met (RSV^R18M^, RSV^R44M^, and RSV^K62M^) to test decreased nucleic acid binding contributing to infectivity loss in RSV^K36M^. Statistics: N=4-5; NS Not Significant, * p<0.05, ** p<0.01; pairwise Wilcoxson rank sum test.

Lys and other positively charged amino acids in NC contribute to nucleic acid binding, Gag multimerization, and binding dynamics at the plasma membrane (34, 38–44). To address if the loss of the Lys in RSV^K36E/M^ contributes to dysfunction because of the lost NLS and not simply to a lost basic residue (42–44), we engineered three other point mutations in NC where an amino acid with a basic charge was changed to Met (NC-R18M, -R44M, and -K62M) and tested function as described above. All three mutants displayed reduced spread through tissue culture compared to RSV^WT^ on day six with RSV^R18M^ spreading to 60%, RSV^R44M^ to 72%, and RSV^K62M^ to 65% (N=4-5, p<0.05, pairwise Wilcoxson rank sum test, Fig 4B). The reduction in spread with these three mutants was as severe as with RSV^K36M^ (N=4-5, NS, pairwise Wilcoxson rank sum test, Fig 4B) and is consistent with a deficiency in nucleic acid binding in contrast to the inability to cycle through the nucleus.

### Replication competent RSV^K36M^ virus does not regain the ability to cycle

Multiple passages through tissue culture inherently increase the potential for rescue mutants to arise. Mutant phenotypes, such as deficient nuclear cycling, can be rescued through reversion of the mutation, which in the case of a single point mutation in RSV^K36M^ is likely and could potentially explain the ability of the mutant to infect cells. Additionally, secondary mutations may arise to counter the phenotype produced from the original mutation.

Furthermore, RSV^K36M^ may localize Gag differently in the context of full-length virus since cellular localization of the RSV-Gag^K36M^ mutation was tested in a limited construct consisting of a truncated Gag-GFP fusion protein. To address the discussed possibilities and address the overall phenotype of full-length RSV^K36M^, we performed immuno-fluorescence staining against RSV-CA to visualize Gag localization. Briefly, DF1 cells infected with replication competent virus from Fig 4 were plated on glass coverslips, treated with or without LMB (10 ng/mL) for 1 hr, and fixed in 5% PFA. Samples were blocked, probed with RbαRSV-CA, probed with fluorescently labeled GtαRb, and nuclei stained with Hoechst. RSV-CA localization in relation to Hoechst-stained nuclei was then visualized under confocal-microscopy. At steady state without LMB treatment, both RSV^WT^ and RSV^K36M^ was expressed in the cytoplasm with no Hoechst co-localization (Fig 5A, WT-left and K36M-left columns). After LMB (10 ng/mL) treatment for 1 hr, RSV^WT^ was expressed in the nucleus with Hoechst co-localization and noticeably less expression in the cytoplasm (Fig 5A, WT-right column). In contrast, close inspection of LMB treated RSV^K36M^ infected cells showed a small amount of Gag nuclear localization compared to untreated cells; however, the majority of the stain remained cytoplasmic with no Hoechst co-localization (Fig 5A, K36M-right column). Localization of Gag in RSV^R18M^, RSV^R44M^, and RSV^K62M^ infected cells displayed primarily nuclear accumulation with a minor deficiency compared to RSV^WT^ (Fig 5B). Together, these data point to a lack of a rescued nuclear cycling phenotype in RSV^K36M^ and suggest that nuclear cycling is not required for RSV infectivity.

**Figure 5.**
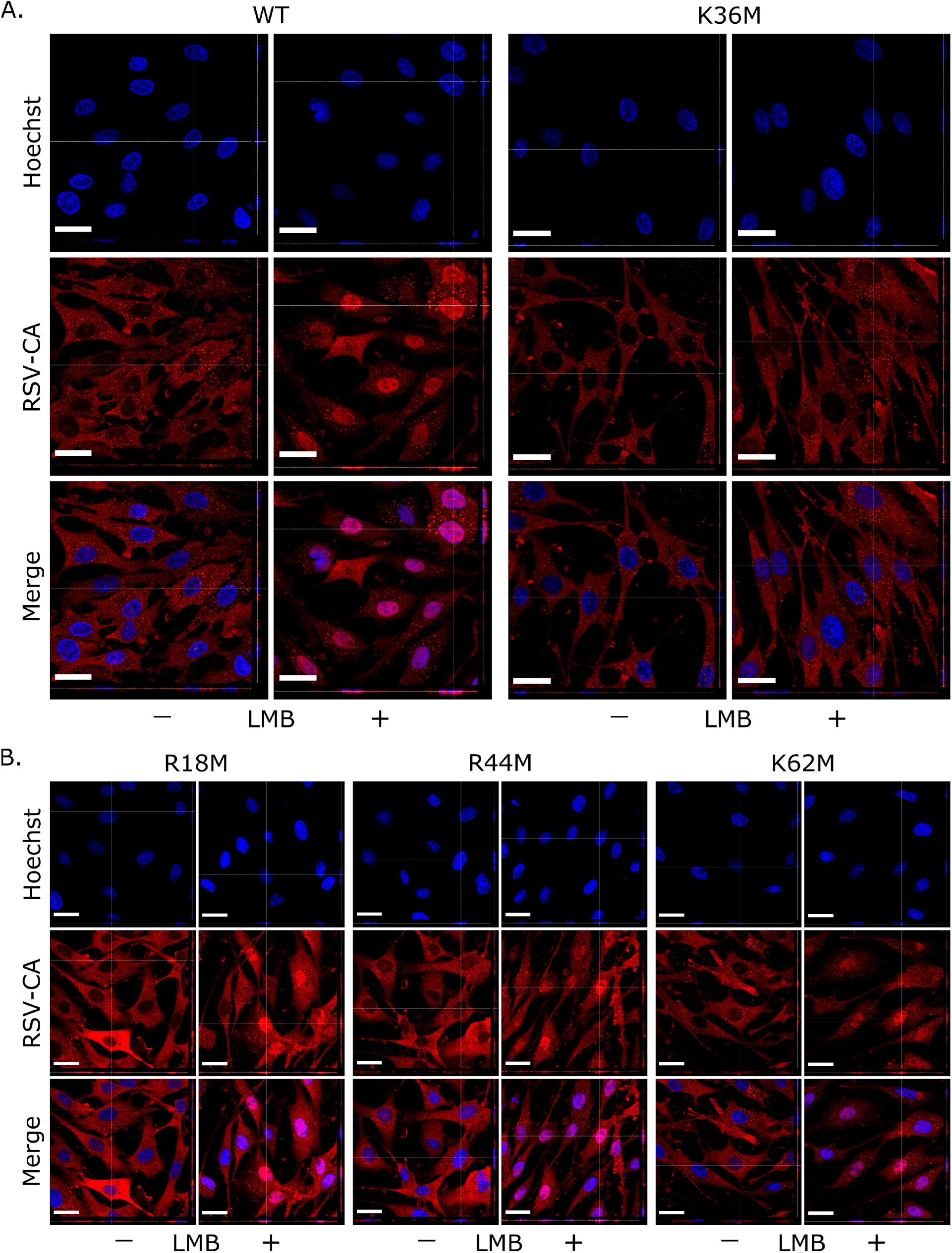
Replication competent RSV^K36M^ is defective in nuclear cycling. Immunofluorescence stains against RSV-CA at steady state. Briefly, DF1 cells infected with replication competent virus from Fig 4 were plated on glass coverslips, treated with or without LMB (10 ng/mL) for 1 hr, and fixed in 5% PFA. Samples were blocked, probed with RbαRSV-CA, probed with fluorescently labeled GtαRb, and nuclei stained with Hoechst. After mounting on slides, cells were imaged with a confocal microscope. (A-B) Representative images and z-stacks of (A) RSV^WT^ and RSV^K36M^ virus as well as (B) RSV^R18M^, RSV^R44M^, and RSV^K62M^ virus with and without LMB treatment. Scale Bar: 20 µm.

### Genomic RNA incorporation in RSV^K36M^ virus does not statistically differ from RSV^WT^

As discussed in the introduction, Gag nuclear cycling was speculated to be the trans-acting mechanism for guiding unspliced gRNA out of the nucleus for subsequent packaging into virions (18–21). Infectivity loss of the previously described mutants was attributed to a loss in gRNA virion incorporation (18, 20, 21). To address whether loss in gRNA virion incorporation contributed to the decreased rate of spread of RSV^K36M^, we quantified gRNA in virions from four independent experiments. Briefly, RSV^WT^, RSV^K36M^, RSV^R18M^, RSV^R44M^, and RSV^K62M^ infected as well as uninfected DF1s at the endpoint of the replication competent experiment (Fig 4) were plated equally and equal amounts of media were collected and concentrated over a 20% sucrose gradient. Equivalent volumes of concentrated media from the four samples were used in parallel for immuno-blot against RSV-CA and qPCR.

Western blots show that relatively equal amounts of virus were produced from the infected cells and were collected for each of the mutants compared to WT. Quantified RSV-CA levels were not statistically different except between RSV^WT^ and RSV^K62M^ (N=4, p<0.05, pairwise Wilcoxson rank sum test, Fig 6A). To quantify genome incorporation, we amplified a 172-base pair segment flanking the NC-NLS region in triplicate for each of four of the five independent samples from Fig 4B. To address potential effects of improper endogenous reverse transcriptase (RT) extension of genome in the RSV mutants due to improper gRNA binding by NC, we performed the reverse transcription with exogenous Moloney MLV in addition to the endogenous RT. We observed a small non-statistically significant decrease in RSV^K36M^, RSV^R18M^, RSV^R44M^, and RSV^K62M^ gRNA incorporation compared to RSV^WT^ (N=3-4, * p<0.05, pairwise Wilcoxson rank sum test, Fig. 6B). The observe non-significant trend may likely need more sensitive assays to parse. Overall, these data suggest that per mL of media relatively equal amounts of virus were produced with relatively equal amounts of genome incorporated in them. Furthermore, these data point toward RSV^K36M^ playing a dysfunctional role in a different portion of the viral life cycle and Gag nuclear cycling serving to enhance efficiency of viral spread.

**Figure 6.**
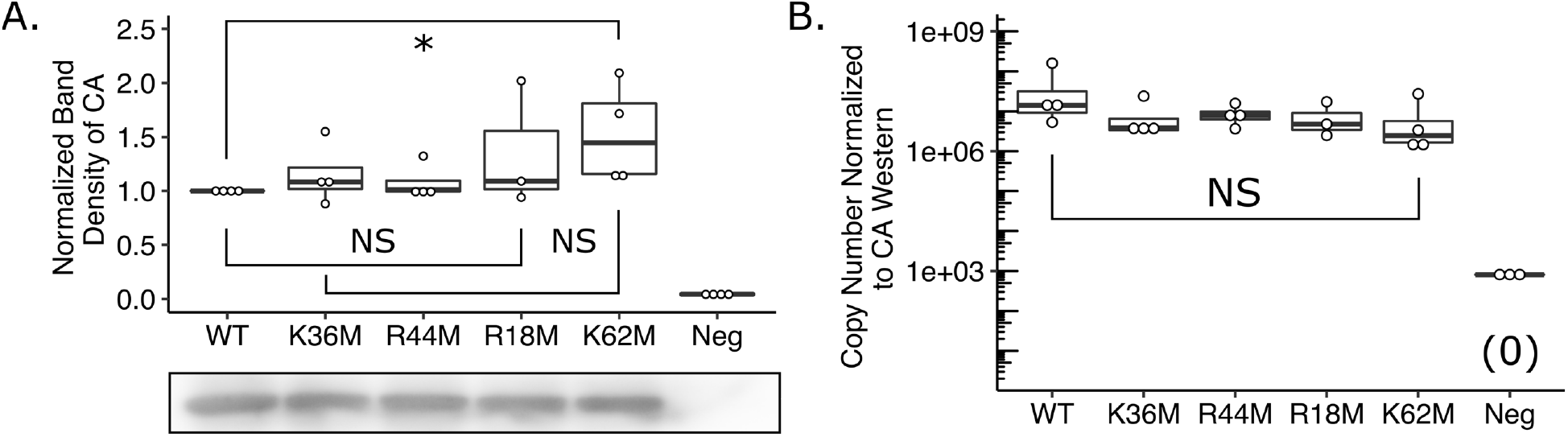
Genomic RNA incorporation in viral particles does not differ between WT and mutants. Infected cells at the end of the replication competent experiment (Fig 4) were plated equally and equal amounts of media were collected and concentrated over a 20% sucrose gradient. Equivalent volumes of concentrated media from the four samples were used in parallel for immuno-blot against RSV-CA and qPCR. (A) Quantification and representative western blot against RSV-CA. (B) Copy number per mL of pre-cleared media normalized to ratios of RSV-CA from western quantification (panel A). After reverse transcription, viral cDNA was amplified flanking NC and mutations were confirmed via sequencing (data not shown). Statistics: N=3-4; NS Not Significant, * p<0.05; pairwise Wilcoxson rank sum test.

## Discussion

While the underlying mechanisms of Gag protein membrane binding and subsequent escape of virions are being established and further refined, intermediate trafficking of Gag proteins and association with gRNA remains elusive and difficult to test. This, in part, is due to the difficulty of tracking individual proteins and other viral components from production to incorporation into virions. When coupled with the multifunctional nature of viral proteins where the same domain can be involved with many aspects across the whole viral lifecycle, this problem is compounded (45, 46). For example, retroviral NC is involved with nucleic acid binding during the late phase by binding and ensuring gRNA incorporation which also results in boosting Gag multimerization (42, 47, 48). During the early phase, NC binds the tRNA^Lys^ necessary for reverse transcription and aids in protecting gRNA/DNA in transit to the nucleus (49). In the case of RSV, the Gag protein rapidly cycles into and out of the nucleus via NLSs in MA and NC and an NES in p10 (9–16). While trafficking of gRNA out of the nucleus for subsequent virion packaging has been implicated as the function of Gag nuclear cycling (9, 10, 18, 21), additional functions and requirement remained to be determined.

### Identifying how HIV-MA blocks nuclear cycling of H/RSV-Gag^WT^

The role of RSV-Gag^WT^ nuclear cycling was first questioned upon observation that an H/RSV-Gag^WT^ chimera where RSV-MA is replaced with HIV-MA did not cycle through the nucleus but remained infectious (28). Previously described RSV non-nuclear cycling mutants attributed the phenotype to strong plasma membrane targeting overcoming the NLS (12, 20). The resulting non-infectious mutants had severe reductions in genome incorporation and both the lack of infectivity and genome was attributed to the lack of nuclear cycling. However, plasma membrane targeting resulted in increased budding efficiency and may have packaging consequences for the virus separate from abolished nuclear cycling. By strongly targeting to the plasma membrane, the virion likely assembles too quickly for proper incorporation of necessary viral components like genome.

We hypothesized that HIV-MA myristoylation provided the more moderate plasma membrane targeting resulting in the abolished nuclear cycling of H/RSV-Gag^WT^. However, introducing a known mutation (G2A) that abolishes myristoylation did not recover nuclear cycling (35). Truncation of HIV-MA in an attempt to identify the reason for abolished nuclear cycling revealed the first α-helix and basic residues from the N-terminal end of the second α-helix to be important for blocking nuclear cycling. Close examination of the crystal structure of HIV-MA shows that the first and second α-helices form a protrusion of basic residues (36). Charged residues play many roles in cellular and viral functions. The truncation of the basic residue protrusion likely causes a conformational change that masks the NC-NLS or removes a strong targeting motif counteracting the NC-NLS of RSV-Gag^WT^ (Fig 1B), though more experiments would be needed to probe the mechanism. H/RSV-Gag^H3^ likely displays cytoplasmic steady state expression without LMB addition due to the removal of acidic residues in helix-2 that mask the basic residues of helix-5 (Fig 1B). Further removal of helix-3 acidic residues in H/RSV-Gag^H5^ likely fully un-masks the basic residues of helix-5, allowing for stronger steady state cytoplasmic retention without LMB addition (Fig 1B).

### Separating plasma membrane trafficking from nuclear cycling with a minimal mutation

A single-cycle chimera is not representative of WT virus, so we sought a minimal RSV mutant that separated plasma membrane targeting from nuclear cycling. We, thus, redirected our efforts to the more centrally located NC-NLS, which would theoretically separate plasma membrane binding from nuclear cycling. A polymorphism at position 36 of NC of RSV strain JS11C1, is defective in Gag nuclear cycling. Distribution of the polymorphism reconstituted in RSV-Gag^K36E^ as well as the less drastic RSV-Gag^K36M^ mutation is diffuse throughout the cytoplasm, similar to WT virus and in constrast to the strong plasma membrane bound phenotype of previously described non-nuclear cycling mutants. Furthermore, RSV^K36M^ remains replication competent, albeit with modestly reduced rate of viral spread. Together, these data suggest that plasma membrane trafficking and binding of RSV^K36M^ is similar to RSV^WT^, albeit the defective nuclear cycling.

The reduced rate of spread does not appear to be due to reduced genome incorporation between RSV^WT^ and RSV^K36M^. Basic amino acid residues are important for nucleic acid binding (42–44). However, single basic amino acid to Met substitutions at other positions of NC neither affected nuclear cycling nor reduced genome incorporation compared to WT, but reduced overall rate of spread to the same level as RSV^K36M^. This points toward an undetectable (with the assays used here) gRNA binding deficiency from the lack of Gag nuclear cycling rather than overall loss of gRNA packaging.

While it could be argued that even small amounts of Gag nuclear localization (as seen in Fig 5A) is enough to traffic gRNA out of the nucleus, one would also expect a more severe loss in rate of spread with a mutation in the NLS as compared to basic residue to Met mutations not in the NLS. The rate of spread is similarly reduced in all NC mutants tested compared to RSV^WT^ (Fig 4B), nuclear localization was not abolished in RSV^R18M^, RSV^R44M^, or RSV^K62M^ (Fig 5B), and genome was incorporated similarly between all mutants (Fig 6B). Together, these pieces of evidence suggest that the dysfunction in replication lies in nucleic acid binding and not the ability to cycle through the nucleus.

## Conclusions

Here, we show that the distinct nuclear cycling that is characteristic of RSV Gag is not required for viral infectivity. However, RSV Gag nuclear cycling is an intriguing phenotype whose function remains elusive. In contrast to “complex” retroviruses like HIV, RSV seems to rely on cis-acting elements on the genome in conjunction with host cellular components to export the genome out of the nucleus similar to other “simple” retroviruses. Gag nuclear cycling, however, boosts rate of viral spread and may be a result of more effective gRNA-virion assembly. Further investigation to identify function may reveal insights to viral protein trafficking and host-virus interactions such as the evolution of trans-acting elements for genome export.

## Materials and Methods

### Plasmid constructs

RSV Gag.3h-GFP (John Wills, Pennsylvania State University) contains full length Gag with PR removed and fused to GFP and will be referred to as RSV-Gag^WT^. Gag.3h-GFP^L219A^ was previously described and will be referred to as RSV-Gag^L219A^ (13). RSV-Gag^ΔMA^ was generated first by PCR amplification of RSV-Gag^WT^ between the start of p10 and the p10 FseI site to engineer an Xhol I site and start codon directly upstream of p10. Secondly, we used restriction digest (NEB restriction enzymes) to remove the region between XhoI upstream of MA and FseI. Products from these two reactions were ligated via In-Fusion HD Cloning Kit (Clontech^®^, 639650). RSV-Gag^ΔNC^ (Volker Vogt, Cornell University) is Gag with NC and PR removed and fused to GFP. RSV-CANC was engineered by PCR amplification of the CA region of RCAS-GFP (Stephen Hughes, NCI-Frederick) to introduce a SacI site followed by a start codon at the 5’ end leading to the SbfI site in the middle of CA. RSV-Gag^WT^ was then digested with SacI and SbfI and ligated to the amplified piece. RSV-Gag^K36E/M^ was engineered into RSV-Gag^WT^ and RSV-Gag^L219A^ by restriction digest to remove the region between SbfI at CA and PspOMI at EGFP with subsequent ligation of PCR products via In-Fusion HD Cloning.

H/RSV-Gag^WT^ has been previously described (28). Truncation mutants were generated by PCR amplification of H/RSV-Gag^WT^ with a forward primer that started at the SacI restriction site, spanned the start Met, and bridged to the helix of interest with a reverse primer from the FseI restriction site. H/RSV-Gag^WT^ was then digested with SacI and FseI and products were ligated via In-Fusion HD Cloning. The resulting construct thus contained the starting Met followed by the helices of interest.

RCAS-GFP was used for replication competent RSV^WT^. RSV^K36E/M^ was engineered by two-step PCR amplification between SbfI at CA and SnaBI at RT with site-directed mutagenesis of K36, restriction digest to remove the region between the two sites, and ligation of products via In-Fusion HD Cloning. NC-R18M, -R44M, and -K62M mutations were engineered into RSV^WT^ by two-step PCR amplification between SacII in CA and SnaBI in RT with site directed mutagenesis at the referenced amino acid positions, restriction digest to remove the region between the two restriction sites, and ligation of products via In-Fusion HD Cloning. RSV^ΔEnv^ in the two-color single-cycle provirus system has been previously described (50). Briefly, Env was removed from RSV^WT^ and was further modified to contain a second reporter—in this case Crimson—outside of the downstream LTR. RSV^ΔEnv,K36E/M^ was generated by restriction digest of RSV^ΔEnv^ and RSV^K36E/M^ at SacII at CA and AgeI at IN, and ligation of products. The two-color reporter system is pseudotyped with VSV-g (NIH AIDS Reagent Program) on a separate plasmid (51).

### Cells

The DF1 cell line was obtained (ATCC, CRL-12203) and maintained in Dulbecco’s modified Eagle’s medium (DMEM, Sigma, D6429-500ML) supplemented with 7.5% fetal bovine serum (FBS, Gibco, 10437-028), 1% chicken serum (Sigma, C5405), 2 mM L-glutamine (Sigma, G7513-100ML), 1 mM sodium pyruvate (Sigma, S8636-100ML), 10 mM minimal essential medium nonessential amino acids (Sigma, M7145-100ML), and 1% minimal essential medium vitamins (Sigma, M6895-100ML). DF1s stably infected with RSV^WT^ and derived mutants were similarly maintained.

### Virus production

RSV VLPs were produced by FuGENE® 6 Transfection Reagent (Promega, E2691) transfection of DF1s at 50% confluence with 1 µg of viral plasmid. For single-cycle virus, VSV-g was added in a 1:9 ratio. Media containing virus (Viral Media) was collected two days post transfection by aspiration. Viral media was then frozen at −80°C for a minimum of 1 hr to lyse cells, thawed in a 37°C water bath, precleared by centrifugation at 3000 x g for 5 min, and supernatant collected by aspiration. Aliquots were stored at −80°C and subsequently used for assays. For assays requiring viral concentration, supernatant collected after preclearing was pelleted through a 100 µL 20% sucrose cushion (20% sucrose, PBS) for 2 h at 30000 x g at 4°C. Supernatant and sucrose buffer was aspirated off leaving a small amount (∼10 µL) of sucrose buffer so as not to aspirate the viral pellet. Viral pellets were stored at −80°C.

### Infectivity assays and flow cytometry

For single-cycle infectivity assays, DF1 cells were transfected at 50% confluency in 6-well format. Five days post transfection, all cells were collected for flow cytometry. For replication competent infectivity assays, DF1 cells at 50% confluency were transfected in 6-well format. To remove the variable of both transfected and infected cells in transfected culture, viral media was collected five days post transfection. Fresh cells were then plated in separate 60 mm dishes and transduced with 500uL of viral media. Cells were then passaged till the population of GFP (+) cells approached 100% as determined by flow cytometry. Infected and non-infected cells were then co-cultured starting at 1% infected. Half of the cell population was collected every two days for flow cytometry.

Cells were collected by washing with PBS and treated with 10 mM TrypLE™ Express Enzyme (Gibco; Cat. No. 12605028). Lifted cells were then collected with PBS and added to 10% PFA to a final concentration of 5%. After 10–20 min incubation at room temperature, the cells were centrifuged at 300 x g for 5 min, supernatant removed, and 300 µL of PBS added. Cells were analyzed for fluorescence using a BD Accuri C6 flow cytometer.

### Microscopy

Fluorescence microscopy was performed on an Olympus IX70 inverted microscope using a Qimaging Rolera Fast camera and Qimaging software. Cells were first plated on glass bottom dishes (MatTek, P35G-1.5-14-C) and transfected with the afore mentioned plasmids. 16-18 hrs post transfection cells were imaged followed by LMB treatment and subsequent imaging. Images were captured using a 100x oil immersion objective. Gain was adjusted so as not to over-expose cells in focus. Confocal fluorescence microscopy was performed on a Leica TCS SP8 inverted spectral confocal microscope. Cells from the replication competent assay were first plated on glass coverslips in 6-well format. 24 hrs post plating, cells were fixed with 5% PFA. Cells were blocked in 5% goat serum in PBST (10% Tween-20) for 1 hr at room temperature, followed by incubation in RSV-CA antibody raised in rabbit (NCI-Frederick, NCI 8/96) at a 1:1000 dilution in blocking buffer. Cells were then washed with PBST (10% Tween-20) 3 x 5min, followed by incubation with GtαRabbit AlexaFluor 555 at a 1:10000 dilution in blocking buffer. Cells were washed once with PBST and 1uL of Hoechst, followed by 3 x 5 min washes with PBST. Cover slips were then mounted on microscope slides, sealed with clear nail polish, and stored at 4°C. Images were captured using a 100x oil immersion objective and z-stacks were captured as optimized by Leica LASX software.

### Western Blot

One mL of supernatant collected after preclearing thawed media containing virus was pelleted via centrifugation through a 20% sucrose cushion (20% sucrose, PBS) for 2 h at 30000 x g at 4°C. Supernatant and sucrose buffer were aspirated off, leaving a small amount of sucrose buffer so as not to aspirate the viral pellet (∼10 µL). Ten µL of 2x sample buffer (50 mM Tris, 2% sodium dodecyl sulfate [SDS], 20% glycerol, 5% β-mercaptoethanol) was added to pelleted virus and heated to 95°C for 5 min before loading.

Cell samples were washed with PBS and trypsinized with 10 mM EDTA. Cells were then collected with PBS, centrifuged at 300 x g for 5 min, and supernatant removed. Twenty µL of RIPA extraction buffer with protease inhibitor was then added to each sample (52). The samples were then kept on ice and vortexed every 5 min for 20 min, followed by centrifugation at 10000 rpm for 10 min at 4 °C. Supernatant was then transferred to a new tube, 20 µL of 2x sample buffer added, and heated to 95 °C for 5 min before loading.

Samples were separated on a 10% SDS-PAGE gel and transferred onto a 0.22 µm pore size polyvinylidene difluoride (PVDF) membrane. Membranes were blocked for 1 hr at room temperature with 4% nonfat dry milk in PBST (10% Tween-20). Membranes were then incubated with anti-RSV-CA antibody raised in rabbit (NCI-Frederick, NCI 8/96) at a 1:500 dilution in blocking solution for 1 hr at room temperature. After blots were washed with PBST (3 x for 5 min), a goat anti-rabbit peroxidase (HRP)-conjugated secondary antibody (Sigma, A0545) was applied at 1:10,000 dilution in blocking solution. After 1 hr, blots were again washed 3x with PBST and imaged. Immobilon Classico Western HRP substrate (Millipore) was used for visualization of the membranes with a chemiluminescence image analyzer (UVP BioSpectrum 815 Imaging System).

### Genome incorporation via qPCR

One mL of supernatant collected after preclearing thawed media containing virus was pelleted via centrifugation through a 100 µL 20% sucrose cushion (20% sucrose, PBS) for 2 h at 30000 x g at 4°C. Supernatant and sucrose buffer were aspirated off, leaving a small amount of sucrose buffer so as not to aspirate the viral pellet (∼10 µL). Sample (∼5 µL) was reverse transcribed with added Moloney MLV. Virus with 0.5 µL each of 100 µM Oligo-dT and 100 µM random hexamers suspended to 6.5 µL dH_2_O was incubated at 65°C for five min followed by ice bath. Clonetech SMART MMLV RT 5xFirst Strand buffer, 10 mM dNTP mix 100 mM DTT, and SMART MMLV RT was then added as per the manufacturer’s protocol. Samples were then incubated at 42°C for 2 hrs followed by 85°C for five min to heat kill the RT.

Reverse transcribed cDNA was then carried forward for qPCR via a BioRad CFX-Connect Real-Time PCR thermal-cycler. 3 µL of cDNA was used in conjunction with the BioRad iTaq Universal SYBR Green Supermix as per the manufacturer’s protocol for 10 µL reactions in a semi-hard qPCR 96-well plate. Each sample was plated in triplicate on the 96-well plate. Known dilutions of RSV^WT^ plasmid were used for standard curve calculation. Using the CFX Maestro software, data was exported as an Excel spreadsheet.

### Data analysis

Flow cytometry data was analyzed using FlowJoTM software (53). Values for fluorescence were exported to Excel spreadsheet. Images were analyzed using Fiji (ImageJ) (54). Western blot images were converted to 8-bit, Fiji’s gel analysis tools used to calculate density, and values exported to an Excel spreadsheet. qPCR data was outputted into Excel spreadsheets. Excel spreadsheets were formatted for statistical analysis via R (55) and exported to CSV format. RStudio was used to analyze data and create figures (56). UGene was used for plasmid cloning, sequence analysis, and multiple sequence alignment image generation (57). Final figures were prepared using Inkscape (58).

## Acknowledgments

We thank the University of Missouri DNA Core Facility for sequencing support. We also thank the University of Missouri Molecular Cytology Core facility for confocal microscopy support. This work was supported by the National Institute of Allergy and Infectious Diseases (NIAID; https://www.niaid.nih.gov) under award R21AI143363 to MCJ.

